# The coalescent for prokaryotes with homologous recombination from external source

**DOI:** 10.1101/151308

**Authors:** Tetsuya Akita, Shohei Takuno, Hideki Innan

## Abstract

The coalescent process for prokaryote species is theoretically considered. Prokaryotes undergo homologous recombination not only with other individuals within the same species (intra-specific recombination) but also with other species (inter-specific recombination). This work particularly focuses the latter because the former has been well incorporated in the framework of the coalescent. We here developed a simulation framework for generating patterns of SNPs (single nucleotide polymorphisms) allowing integration of external DNA out of the focal species, and a simulator named **msPro** was developed. We found that the joint work of intra- and inter-specific recombination creates a complex pattern of SNPs. The direct effect of inter-specific recombination is to increase the amount of polymorphism. Because inter-specific recombination is very rare in general, it creates a regions with an exceptionally high level of polymorphisms. Following an inter-specific recombination event, intra-specific recombination chop the integrated foreign DNA into small pieces, making a complicated pattern of SNPs that looks as if foreign DNAs were integrated multiple times. This work with the **msPro** simulator would be useful to understand and evaluate the relative contribution of intra- and inter specific recombination to creating complicated patterns of SNPs in prokaryotes.

---

The coalescent is a population genetic theory, which considers the evolutionary process backward in time (Kingman 1982; Hudson 1983b; Tajima 1983). The coalescent theory has been mainly developed by assuming its application to higher eukaryotes, perhaps due to a historical reasons: The major model species of population genetics have been higher eukaryotes such as Drosophila and human (*e.g.,* Hartl and Clark 2007). The coalescent provides an extremely powerful simulation tool for analyzing the pattern of single nucleotide polymorphisms (SNPs) in sampled sequences. It is flexible enough to incorporate major evolutionary processes including random genetic drift, mutation, recombination and demographic history (*e.g.,* Hudson 1990; Nordborg 2001; Wakeley 2008), whereas it is not very straightforward to incorporate complex modes of selection (but see Krone and Neuhauser 1997; Neuhauser and Krone 1997; Donnelly and Kurtz 1999; Fearnhead 2006). **ms** is one of the most popular coalescent simulators, which allows to produce patterns of neutral SNPs under various settings of demography (Hudson 2002). It incorporates two major outcomes of meiotic recombination, that is, meiotic crossing-over and gene conversion.

Prokaryotes are unique in that they are haploids and do not undergo meiosis, and therefore their recombination mechanisms are quite different from that of meiotic recombination in eukaryotes. Nevertheless, the coalescent can work with prokaryotes with a relatively simple modification: Recombination is treated as an event analogous to meiotic gene conversion because a prokaryote’s circular chromosome needs double “crossing-over” to exchange a DNA fragment. This modification can well explain the nature of prokaryotes’ homologous recombination as we will explain below. The application of the coalescent theory to bacteria became particularly popular since McVean *et al.* developed the software **LDhat** (McVean *et al.* 2002) for estimating the recombination rate, which is a modified version of Hudosn’s composite likelihood method (Hudson 2001). Because **LDhat** allows recurrent mutations at a single site, it is more suitable to species with a large population size like bacteria. **LDhat** has been applied to the multilocus sequence typing (MLST) data (*e.g.,* Jolley *et al.* 2005; Pérez-Losada *et al.* 2006; Wirth *et al.* 2006) and genome-wide SNP data from various species (*e.g.,* Touchon *et al.* 2009; Donati *et al.* 2010; Haven *et al.* 2011), demonstrating a great variation in the recombination rate across species. Hudson’s **ms** software has also been successfully used (*e.g.,* Pepperell *et al.* 2010; Thomas *et al.* 2012; Zhang *et al.* 2012; Takuno *et al.* 2012; Cornejo *et al.* 2013; Nell *et al.* 2013; Krause *et al.* 2014; Shapiro 2014; Rosen *et al.* 2015).

Thus, despite the large difference in the recombination mechanism between eukaryotes and prokaryotes, it is technically not very difficult to handle prokaryotes’ homologous recombination in the coalescent framework. However, this holds only when recombination occurs within a single species. This assumption should hold quite strictly in eukaryotes, but not in prokaryotes for which the concept of species is not as strict as eukaryotes (*e.g.,* Cohan 2002b; Doolittle and Papke 2006; Achtman and Wagner 2008) because of frequent exchanges of DNA between different species due to the nature of their recombination mechanism, as is described in the following.

Prokaryotes undergo recombination by incorporating DNA outside of the cell through three major mechanisms: natural transformation, transduction, and conjugation, as illustrated in Figure 1 (*e.g.,* Snyder *et al.* 2013). Natural transformation is a process involving direct uptake of a free extracellular DNA and the integration under natural bacterial growth conditions (Figure 1A). Transduction is a process in which bacterial DNA is introduced into the other bacteria through infection by a phage containing the DNA (Figure 1B). Conjugation is the transfer of DNA from one bacterial cell to another by the transfer functions of a self-transmissible DNA elements, frequently associated with plasmids (Figure 1C).

**Figure 1.**
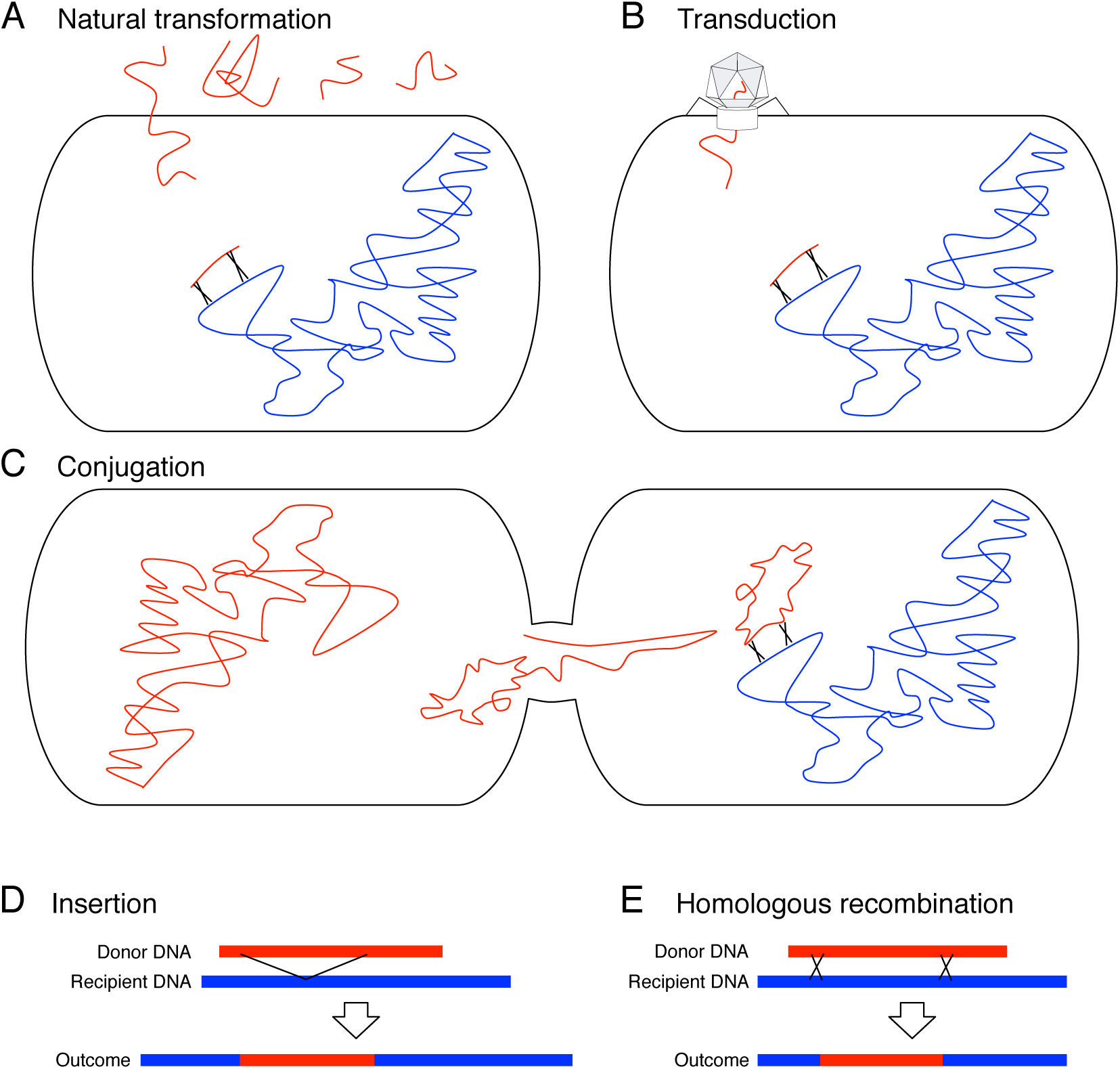
Three major mechanisms for prokaryotes to integrate external DNA: natural transformation (A), transduction (B), and conjugation (C). The host genome and external DNA are presented in blue and red, respectively. (D, E) Two outcomes of recombination via insertion (D) and homologous recombination (E).

It is known that such incorporated DNA from outside of the cell is usually harmful when integrated into the host genome, so that there are a number of mechanisms to avoid integration (*e.g.,* Lorenz and Wackernagel 1994; Majewski 2001; Cohan 2002a; Chen and Dubnau 2004; Thomas and Nielsen 2005; Marraffini and Sontheimer 2010; Vasu and Nagaraja 2013). Because DNAs from different species should be much more harmful than those from the same species, most mechanisms involve some kind of self-recognition systems, in which markers are distributed through the genome to distinguish from those originating external source *(i.e.,* different species). In some bacterial species, such as *Neisseria gonorrhoeae* and *Haemophilus influenzae,* efficient natural transformation requires the presence of short sequence motifs (~10bp), called as DNA uptake sequences (DUS) or uptake signal sequences (USS), that is interspersed among the genome, which may prevent the incoming DNA from different species (or strains) integrating into their genomes. Phage defense mechanisms may also work against incoming DNA from external source. The restriction-modification system is a common mechanism or degrading DNA that is not properly modified (*e.g.,* through DNA methylation), which have been identified in ~90% of prokaryote species (Roberts *et al.* 2010). Clustered, regularly interspaced short palindromic repeat (CRISPR) loci and their associated proteins (Cas) are found in the genomes of ~90% of archaea and ~50% of eubacteria (Grissa *et al.* 2007; Rousseau *et al.* 2009). These sequences are separated by short sequences of DNA (23–50 bp) known as spacers, most of which exhibit homology to previously encountered phage or plasmid genomes, suggesting that these loci provide memory for the bacteria to prevent repeated incoming encounters.

In addition, when these mechanisms do not work perfectly, there is another round of screening process to prevent integration of external DNA through homologous recombination (Majewski 2001). For example, recombination requires near identical regions (*e.g.,* monitored by RecA mediated homology search), (Shen and Huang 1986; Majewski and Cohan 1998) so that external DNA has less chance to be integrated. In addition, it is also pointed out that the mismatch repair system is effective in preventing recombination between highly mismatched sequences (Claverys and Lacks 1986; Majewski 2001; Overballe-Petersen *et al.* 2013). Thus, there are a number of molecular mechanisms to prevent integrating external DNA to the host genome. Nevertheless, it has been repeatedly demonstrated that prokaryote genomes undergo recombination, not only within the same species but also with different species (reviewed in Majewski 2001).

There are two possible outcomes of recombination as shown in Figures 1D and E (see Lawrence 2013, for a review). One is that the incorporated DNA is inserted into the genome (Fig. 1D), and the other is that the incorporate DNA is exchanged with its homologous part of the genome if any (Fig. 1E). The former is known as horizontal gene transfer or lateral gene transfer, and its evolutionary role is emphasized when a novel gene is acquired and contributes to adaptation (Ochman *et al.* 2000; Dobrindt *et al.* 2004; Fraser *et al.* 2009; Polz *et al.* 2013), although the frequency and importance of such illegitimate recombination is under debate (de Vries *et al.* 2001; Shapiro *et al.* 2012). The latter is known as homologous recombination, and it usually involves DNAs from the same species because the near-identity requirement of the RecA mediated homology search criteria is easily satisfied, whereas it is also possible that DNA from different species is integrated as long as it retains some homology. Homologous recombination between different species sometimes remains unique patterns of SNPs, from which we can search for their footprints in the sequence data (reviewed in Awadalla 2003; Didelot and Maiden 2010; Azad and Lawrence 2012; Nakhleh 2013).

The focus of this article is the latter, homologous recombination. Considering the mechanism of homologous recombination involving double crossing-over, the outcome is similar to meiotic gene conversion. Therefore, as mentioned earlier, the standard coalescent has been commonly applied for analyzing patterns of SNPs in bacteria with a simple modification in the setting; the rate of crossing-over is set to zero, so that all recombination events (*i.e.,* homologous recombination) are treated as if they are meiotic gene conversion. This application should be reasonable as long as the donor of homologous recombination is always another individual in the same species. However, it is well known that homologous recombination occasionally involves DNA from other species, and this is the case that the standard coalescent cannot handle. The purpose of this work is to develop the theoretical framework of the coalescent for prokaryotes, which allows homologous recombination both within and between species. We also developed a simulation software named **msPro,** which will be available at http://www.sendou.soken.ac.jp/esb/innan/InnanLab/).

## Theoretical Framework

### Overview

Consider a sample of prokaryote DNA sequences with length *L* bp from *n* haploids, and trace their ancestral lineages backward in time. Figure 2A shows an example of an ancestral recombination graph under the standard coalescent, in which all recombination is assumed to be homologous recombination within the same species (Hudson 1983a; Griffiths and Marjoram 1996). Under this setting, the process is analogous to meiotic gene conversion in the standard coalescent for diploid eukaryotes (McVean *et al.* 2002; Awadalla 2003). A coalescent event merges the ancestral lineages (*e.g.,* events 3, 4, 5, and 6 in Fig. 2A), and homologous recombination separates the lineage into two (*e.g.,* event 1, and 2 in Fig. 2A). For example, event 1 in Fig. 2A is a homologous recombination, in which a short fragment (presented by a gray box) is integrated into the recipient genome, so that the ancestral lineage is separated into two; one for the recipient genome and the other is for the integrated fragment. Then, following the standard treatment, we further trace their ancestral lineages until the lineages of all sampled chromosome merge to their MRCA (Most Recent Common Ancestor), which is referred to as MRCA_all_ in this article. It should be noted that with the presence of recombination, different parts of the region have different histories, so that MRCA_all_ cannot be identical across the region; different subregions chopped by recombination should have their specific MRCA_all._ For example, MRCA_all_ for the black region appears at time *T*_6_, while that for the gray region is at time *T*_5_ and for the other white regions are at *T*_4_ (Fig. 2A). The ancestral recombination graph has all historical information for the entire regions as illustrated in Fig. 2A. With this ancestral recombination graph, a pattern of SNPs can be simulated by randomly distributing point mutations on the graph. Thus, the standard coalescent treatment works for prokaryotes with homologous recombination within species (McVean *et al.* 2002; Awadalla 2003).

**Figure 2.**
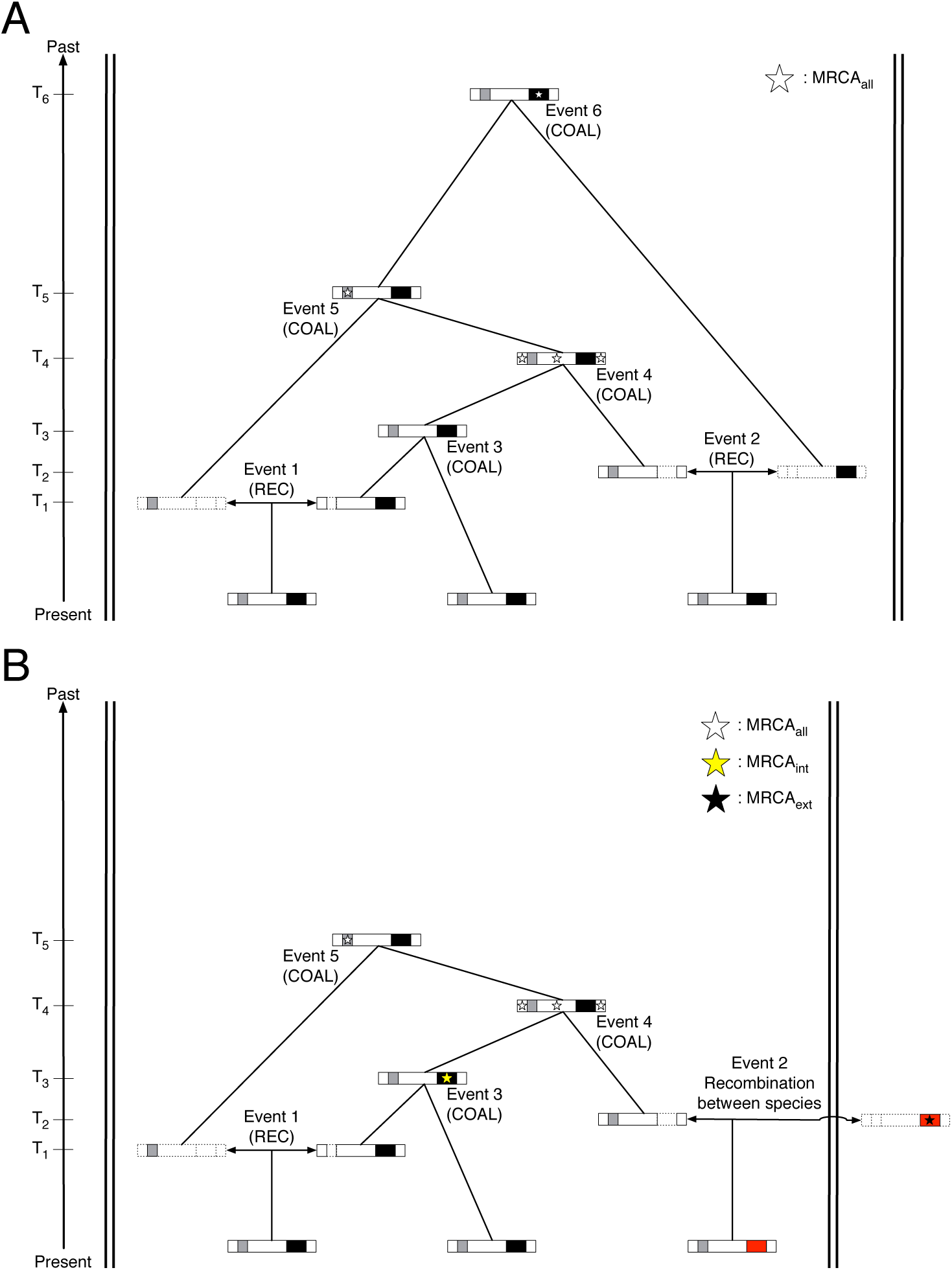
(A) Ancestral recombination graph with recombination within species alone. An example with sample size *n* = 3 is illustrated. The sampled three genomes are shown by long boxes, where regions with different histories are presented in different colors. The ancestral lineage is split into two by a recombination event (REC), while a pair of ancestral lineages merges by a coalescent event (COAL). The boxes with dashed lines represent dummy regions whose descendants do not show up in the sample. The two short regions in gray an black are transferred fragment by gene conversion-like recombination events. MRCA_all_ for each region is shown by a star. The white part has MRCA at *T*_4_, the gray part has MRCA at *T*_5_ and the black part has MRCA at *T*_6_. (B) Ancestral recombination graph with recombination within species and from external source. The region transferred from external source is shown in red.

The problem is when DNA from other species is integrated by homologous recombination. Figure 2B illustrates such a situation, in which event 2 is assumed to be a homologous recombination event from external source (*i.e.,* integration of DNA from other species), which is presented in a red box. In this case, the ancestral lineage of the transferred DNA originates from outside of the focal species, so that it is not involved in the coalescent process of the focal species before time *T*_2_. This is the situation that the standard coalescent cannot handle. We here propose a simple solution to this problem: Tracing the ancestral lineage of external source should be terminated, and the coalescent process should be continued without considering such terminated lineages. Under this treatment, the concept of the MRCA of all sampled sequences (MRCA_all_) does not apply to such a region that experienced homologous recombination from external source. The direct donor of the external DNA is called MRCA_ext,_ most recent common ancestor from external source, and MRCA of the rest is referred to as MRCA_int,_ most recent common ancestor of internal lineages. In event 2 in Figure 2B, while the gray and white regions that are not involved in the integration of external DNA can be traced back to MRCA_all_ (at *T*_5_ and *T*_4_, respectively), we may stop tracing the ancestral lineage of the red region at > *T*_2_, and the origin of this region is treated as a MRCA_ext._ Thus, when a region experienced a homologous recombination from external source, the sampled sequences have two kinds of origins, one is MRCA_ext_ at *T*_2_ as the origin of the red part (shown by a red box with a star in Figure 2B) and the other is MRCA_int_ at *T*_3_ as the origin of the rest, shown by a black box with a yellow star in Figure 2B.

We here consider how to simulate a pattern of SNPs in such a region that experienced homologous recombination with external source. Note that provided the mechanism of homologous recombination, we assume a reasonable level of sequence identity between the external DNA and the focal species. This means that the external lineage should eventually coalesce with the common ancestor of the focal species (on the time scale of species-divergence). However, it is very difficult to know the probability distribution of the time to such eventual common ancestor, which could be far older than the MRCA of the focal species.

Alternatively, we develop an ad-hoc treatment that does not require any unknown ancient demographic history up to species divergence. The idea of our treatment is based on a number of empirical demonstrations that the rate of successful integration of external DNA heavily depends on the nucleotide divergence between the transferred fragment and the recipient sequence; the rate decays almost exponentially with increasing divergence as demonstrated by many authors (Albritton *et al.* 1984; Roberts and Cohan 1993; Vulić *et al.* 1997; Zahrt and Maloy 1997; Lorenz and Sikorski 2000; Majewski *et al.* 2000). See below for details.

### Homologous recombination within species (intra-specific recombination)

It is relatively straightforward to incorporate homologous recombination within species as mentioned above. Following previous studies (Wiuf and Hein 2000; McVean *et al.* 2002), we assume that a homologous recombination event is initiated at any position at rate *g* per site per generation. Then, it is assumed that the elongation process proceeds such that the length of transferred tract, *z,* follows a geometric distribution, with mean tract length = *λ* bp:
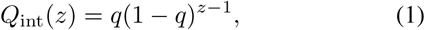

where *q* = 1/λ. This assumption is supported by empirical studies on transformation of many species including *Helicobacter pylori* (Lin *et al.* 2009), *Streptococcus pneumoniae* (Croucher *et al.* 2012), and *Haemophilus influenzae* (Mell *et al.* 2014). For mathematical convenience, we assume unidirectional elongation of conversion tract from 5’ to 3’, which has no quantitative effect on the pattern of SNPs. Given Equation 1, the rate that a region of *L* bp undergoes homologous recombination per generation is given by
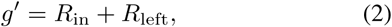

where *R_in_* is the rate of gene conversion initiating inside the region and *R_left_* is the rate outside the region but ending within the observed sequence. *R_in_* and *R_left_* are given by
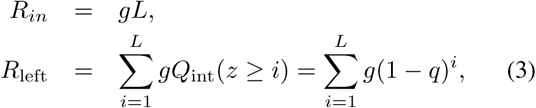

Assuming all recombination is neutral, this rate (*g′*) is identical to the backward recombination rate, which can be directly incorporated into the coalescent framework. The backward recombination rate per generation is defined as the rate at which a lineage undergo recombination when a lineage is traced back for a single generation.

It is interesting to note that Equation 2 does not include the probability that a recombination tract cover the entire simulated region. This is because such recombination simply causes a shift of a lineage to another lineage within the same population, which does not essentially affect the coalescent process. However, this does affect the process if the recombination event occurs with different species as we will explain in the next section (see below).

It should be noted that there are three mechanisms for a cell to incorporate DNA, transformation, transduction and conjugation, through which homologous recombination can occur. They occur at different rates and typical lengths of integrated tracts should be different. Quite short fragments are usually integrated through transformation, while relatively large fragments may be involved in recombination through transduction and conjugation (Cohan 2002a). Therefore, it is biologically reasonable to model these processes separately, and this is what was done in earlier studies (see Maynard Smith 1994; Hudson 1994).

However, in some studies (particularly in coalescent-based studies), all three recombination processes are not specified (*e.g.,* Falush *et al.* 2001; McVean *et al.* 2002; Awadalla 2003; Fearnhead *et al.* 2005). This should be partly because the three mechanisms are commonly summarized by a single backward recombination rate and the tract length is simply assumed to follow a geometric distribution (although not specifically described in these literatures to the best of our knowledge). This may work perhaps because the possible outcome of the three recombination mechanisms are similar in that they can be described as a double-recombination event (Wiuf 2001) even when the typical tract lengths and rates are different (Maynard Smith 1994; Hudson 1994), but we should remember that this is a conventional approximation.

More strictly, if we consider the three mechanisms separately, denote by *g*_tf_, *g*_td_, and *g*_cj_, respectively, the initiation rates of transformation-, transduction- and conjugation-oriented recombination per site per generation, and for each, let us assume that the tract length follows a geometric distribution (the mean lengths are λ_tf_, λ_td_, and λ_cj_ for the three mechanisms). Then, the total initiation rate per site is *g*_total_ = *g*_tf_ + *g*_td_ + *g*_cj_, but the density distribution of tract length is not a simple geometric distribution with a single parameter, rather given by an average of three geometric distributions:
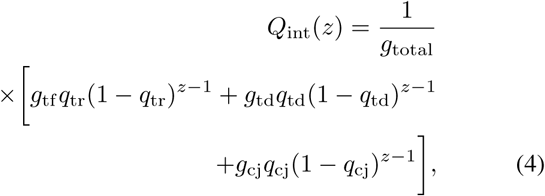

where *q*_tr_ = 1/λ_tr_, *q*_td_ = 1/λ_td_, and *q*_cj_ = 1/λ_cj_. Thus, strictly speaking, there should be situations where the ad-hoc treatment using a single geometric distribution may not hold. Nevertheless, the simplified treatment may work fairly well if we assume that one of the three mechanisms dominates the other two. For example, it is well known that many of *Bacillus* species show a very high transformation rate (especially in laboratory strains of *B. subtilis,* Earl *et al*. 2008), whereas some species are not naturally transformable (*e.g., Escherichia coli, Salmonella typhimurium;* Lorenz and Wackernagel 1994) and conjugation and/or transduction may be should be the major cause of recombination.

Thus, although it is mathematically correct to model the three mechanisms separately, there should be many cases where it is reasonable to use the simplified treatment. This is convenient to apply the coalescent theory to real polymorphism data for estimating the rate of homologous recombination, especially when the relative contributions of the three mechanisms are unknown. In this work, therefore, we employ the simplified treatment with a single rate of homolo gous recombination (Equation 2) with a single geometric distribution with parameter *q* (Equation 1), following previous theoretically studies (Falush *et al*. 2001; McVean *et al*. 2002; Fearnhead *et al*. 2005; Jolley *et al*. 2005; Didelot and Falush 2007).

As mentioned above, a homologous recombination event within prokaryote species is easily incorporated in the standard framework of the coalescent (Wiuf and Hein 2000; McVean *et al*. 2002; Fearnhead *et al*. 2005; Jolley *et al*. 2005). That is, when tracing the ancestral lineage of a certain sequence with *L* bp, the process waits for either coalescent or recombination event, and the per-generation rate for the latter is given by Equation 2, while the rate of coalescence is given by 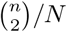, where *N* is the population size and *n* is the number of lineages.

Note that this simple process holds in a single population, in which coalescence occurs randomly between any individuals in the population and so does recombination, but it is straightforward to incorporate population structure and demographic history into this framework as is done for eukaryote cases. The difference between eukaryote and prokaryote is the causes of population structure. In eukaryotes, limited migration between geographic barriers should be the major cause, and this also applies to prokaryotes although more complicated. For example, subpopulations of infectious species may form based on host individuals.

In addition, there are two major classes of isolation in prokaryote, ecological and genetic isolation. Ecological isolation is defined as a difference of niche that can reduce the rate of recombination between bacterial populations (Cohan 2002a,b). Physiological difference between donor and recipient would decrease the chance of recombination. For example, *Vibrio splendidus* lives in coastal bacterioplankton, exhibiting resource partitioning (specific season and/or free-living size fraction) among strains and phylogenetic divergence corresponding to each niche. It is suggested that ecological isolation is working as a barrier of DNA exchanges between niches (Hunt *et al.* 2008). Genetic isolation is defined as the establishment of mutation accumulation that prevents one strain from integrating foreign DNA of other strains (*e.g.,* Lawrence 2013). As described in the Introduction, the rate of successful integration of DNA of other strains depends on a number of self-recognition mechanisms, including short-specific sequences (*i.e.,* DUS or USS), restriction-modification systems, RecA-mediated homology search, and mismatch correction system. While both ecological and genetic isolation are often used in the context of homologous recombination with different species (or strains) involved in sexual isolation (Cohan 2002a; Fraser *et al.* 2009), these concept should work for homologous recombination within the same species but between different populations (or strains). Thus, there are many factors to cause isolation within the same species, which should heavily affect the pattern of the coalescent process. Therefore, when analyzing data with coalescent simulations, past demographic history including such isolations should be well taken into account accordingly (*e.g.,* Kreitman 2000; Nordborg 2001; Rosenberg and Nordborg 2002; Nordborg and Innan 2002; Sousa and Hey 2013).

### Homologous recombination with different species (inter-specific recombination)

We again use the backward argument. We define *h* as the backward recombination initiation rate per site. That is, when tracing the ancestral lineage of a single generation backward in time, *h* is the rate at which the lineage experiences a recombination event from external source that is initiated at the focal site (the same definition as *g* except for the source of integrated DNA). Given *h*, we can compute *h*′, the rate for the simulated region, using a similar equation to (2) (see below for details). With this rate specified, it is very straightforward to incorporate homologous recombination with different species into the coalescent framework: When tracing a lineage of a sequence with *L* bp, the process considers which is the next event, coalescence, recombination within species or recombination from external source, with relative backward rates, 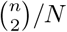, *g*′ and *h*′, respectively. If a recombination event from external source occurs, the length of a transferred region is randomly determined (see below). Then, the transferred region is replaced by a sequence representing external source. Thus, the process can be well merged with the backward treatment of the coalescent, except that the biological interpretation of *h*, the backward rate recombination from external source, should be considered carefully, as we explain in the following.

In order to define *h*, let us consider the coalescent process of a particular species (population), around which there are a number of different species. The focal species potentially undergoes recombination with these species, and the rate of such recombination should be determined by a number of genetic and ecological factors as mentioned above. Figure 3 illustrates a hypothetical situation of a certain species, *S*_0_, around which there are five other species (*S*_1_ − *S*_5_), and their proportion is shown in the pie-chart (Figure 3A). The five species are in the order based on the divergence (d) from *S*_0_. *d* in each species might follow some distribution as illustrated in Figure 3B. Then, the dashed line (lines in five colors combined) in Figure 3B can be considered to represent the density distribution of divergence of DNA sequences that could recombine with the focal species. As mentioned above, the rate of successful integration of these DNA to the focal species varies depending on the species due to the genetic and ecological barriers against recombination. Furthermore, even when recombination successfully occurred, integrated DNA may be deleterious to the host individual and could be immediately selected out of the population. The distribution in the solid line in Figure 3B takes these effects into account, and the degree of reduction for each species is shown by an arrow. Then, noting that the tract length of homologous recombination roughly follows a geometric distribution, we obtain *Q*_ext_ (*d, z*′), the joint distribution of *d* and successfully integrated tract length (*z*′) as illustrated in Figure 3C. The definition of *h* is the per-site rate of such successful recombination from external source. *h* is much smaller than the forward recombination rate because we assume that deleterious recombinations are immediately purged from the population. In other words, we here assume that successfully incorporate foreign DNAs are neutral in the population of the focal species. Under this setting, recombination from external source can be simply incorporated in the coalescent framework as described above: The event of recombination from external source is included at rate *h*′ together with coalescence and recombination within species that occur at rates 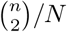 and *g*′, respectively. When recombination from external source occurs, the tract length (*z*′) and nucleotide divergence within the tract (*d*) can be determined as a random variable from *Q*_ext_ (*d, z*′).

**Figure 3.**
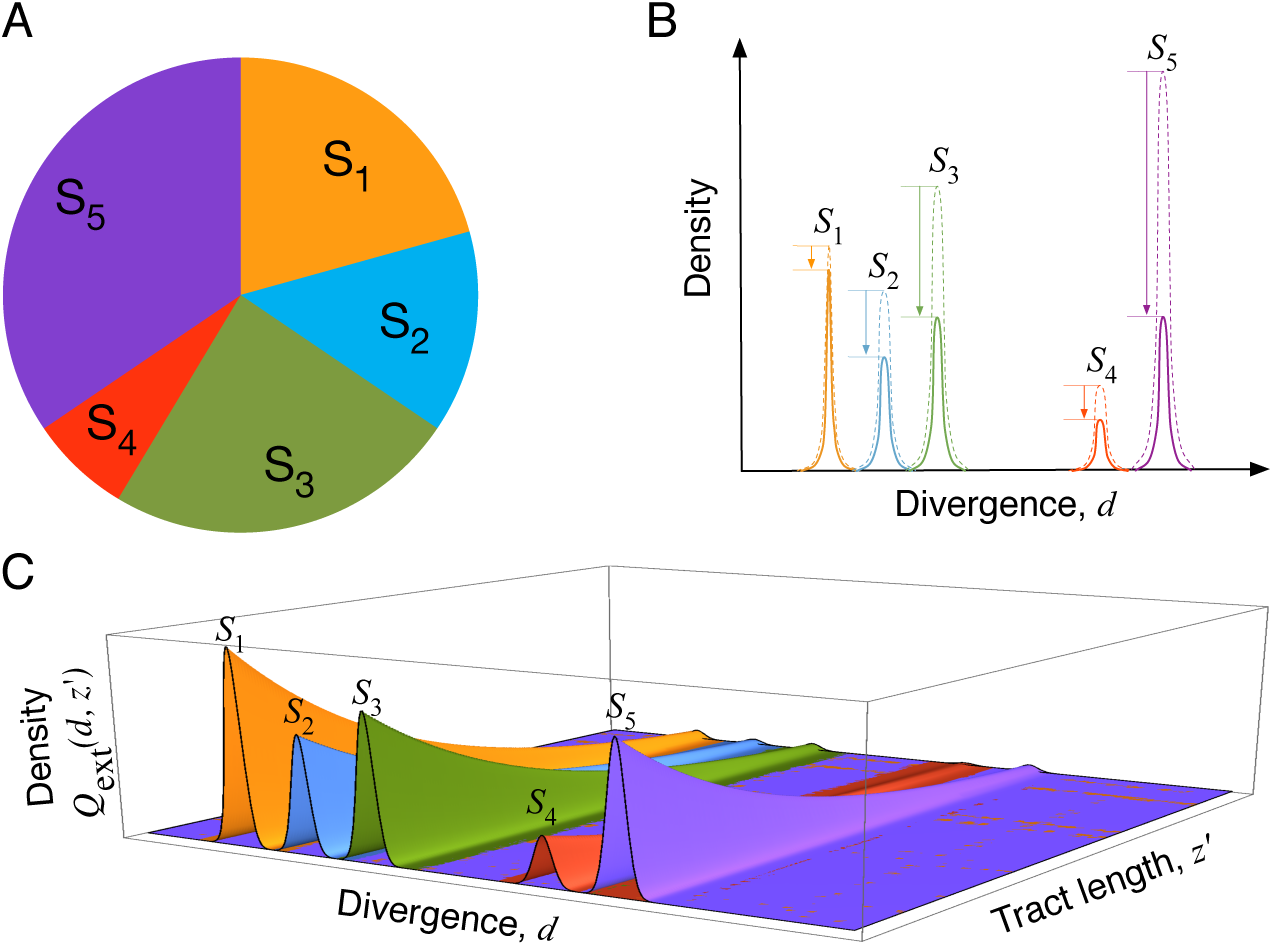
Illustrating a hypothetical environment where five different species (*S*_1_ - *S*_5_) are there around the focal species, *S*_0_. (A) The proportion of the five species in the environment. (B) Density distribution of the divergence of environmental DNA from the focal species (dashed line) and the waited distribution according to the probability of successful integration in the genome of the focal species. (C) The joint density distribution of *d* and successfully integrated tract length (*z*′).

The computation of *h*′ from *h* is slightly different from the treatment for recombination within the same species (see Equation 4), because we cannot ignore the recombination event that encompassed the entire simulated region. That is, *h*′ is given by
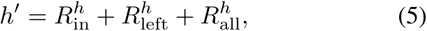

where 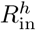 is the rate of recombination initiating inside the region, 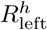 is the rate initiating outside the region and ending within the focal sequence, and 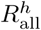 is the rate initiating the 5’ upstream of the region and ending in the 3’ downstream of the region. 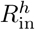 and 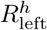 and 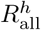 are given by
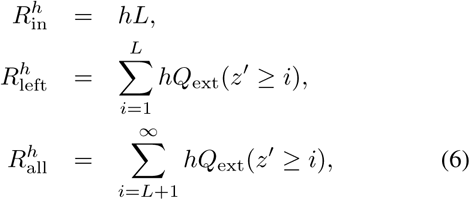

where *Q*_ext_(*z*′) is calculated by integrating the joint probability distribution (*Q*_ext_(*d*, *z*′)) over *d*. From Equations 5-6, *h*′ is written as
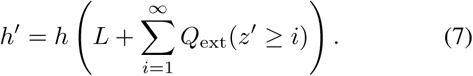

### Mutation

Once an ancestral recombination graph is constructed, neutral point mutations are distributed on it. Our model assumes a finite length of sequence (L bp), and mutation occurs symmetrically between two allelic states at rate *μ* per site per generation, and the population mutation rate is defined as *θ* = 2*N μ.*

## Results and Discussion

We carried out simulations to generate a number of patterns of SNPs to demonstrate the effect of homologous recombination from external source (inter-specific recombination). The mutation rate *θ* = 0.01 was fixed throughout this work. For recombination within the focal species (intraspecific recombination), the mean tract length was fixed to be *λ* = 1000 bp, and the rate (*g*) was changed. We first considered a relatively low recombination rate from external source (2*Nh* = 0.00005). We here used a simplified assumption to demonstrate the point, that is, the average divergence to external DNA was fixed to be 20% and the tract length followed a geometric distribution with a fixed mean *ξ* (*i.e., Q*_ext_(*d* = 0.2, *z*′) = *ξ*^−1^(1−*ξ*^−1^)^*z*′−1^). Figure 4 shows typical patterns of SNPs from the simulation results with *n* = 10, and *L* = 5, 000. The positions of SNPs are presented by solid vertical lines along the simulated region. In Figure 4A, no recombination within species is assumed (2*N g* = 0). One recombination event (607 bp) from external source occurred *t* = 0.23N generations ago on the ancestral lineage of individuals 2, 5 and 10, and the positions of two breakpoints of the recombination event are shown by red arrows. The region that originates from foreign DNA can be clearly recognized as a cluster of SNPs due to large divergence (*d* = 0.2, 20 times larger than *θ*). This region is referred to as Region 1 and boxed in red. Neighbor-jointing tree for this region is completely different form that for the other region: Individuals 2, 5 and 10 are highly diverged from the other seven individuals in Region 1.

It is thus obvious that the level of polymorphism increases as recombination events from external source increases. As shown in Figure 5A, the level of polymorphism increases with increasing the initiation rate (*h*), mean tract length (*ξ*) and divergence (*d*), where the amount of polymorphism is measured by *π*, the average number of nucleotide differences per site. This simulation result agrees with theoretical prediction that the expectation of *π* is given by a simple function of *h, ξ, d:*

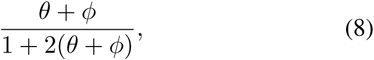

where *ϕ* = 2*Nhξd*. See Appendix for the derivation. It is obvious that the most important parameter is the product of three recombination-associated parameters, *hξd,* which represents the probability that the allelic state at a single site is flipped by recombination (see Appendix), as Figure 5B clearly demonstrates that *π* is given by a simple liner function of *hξd.*

**Figure 4.**
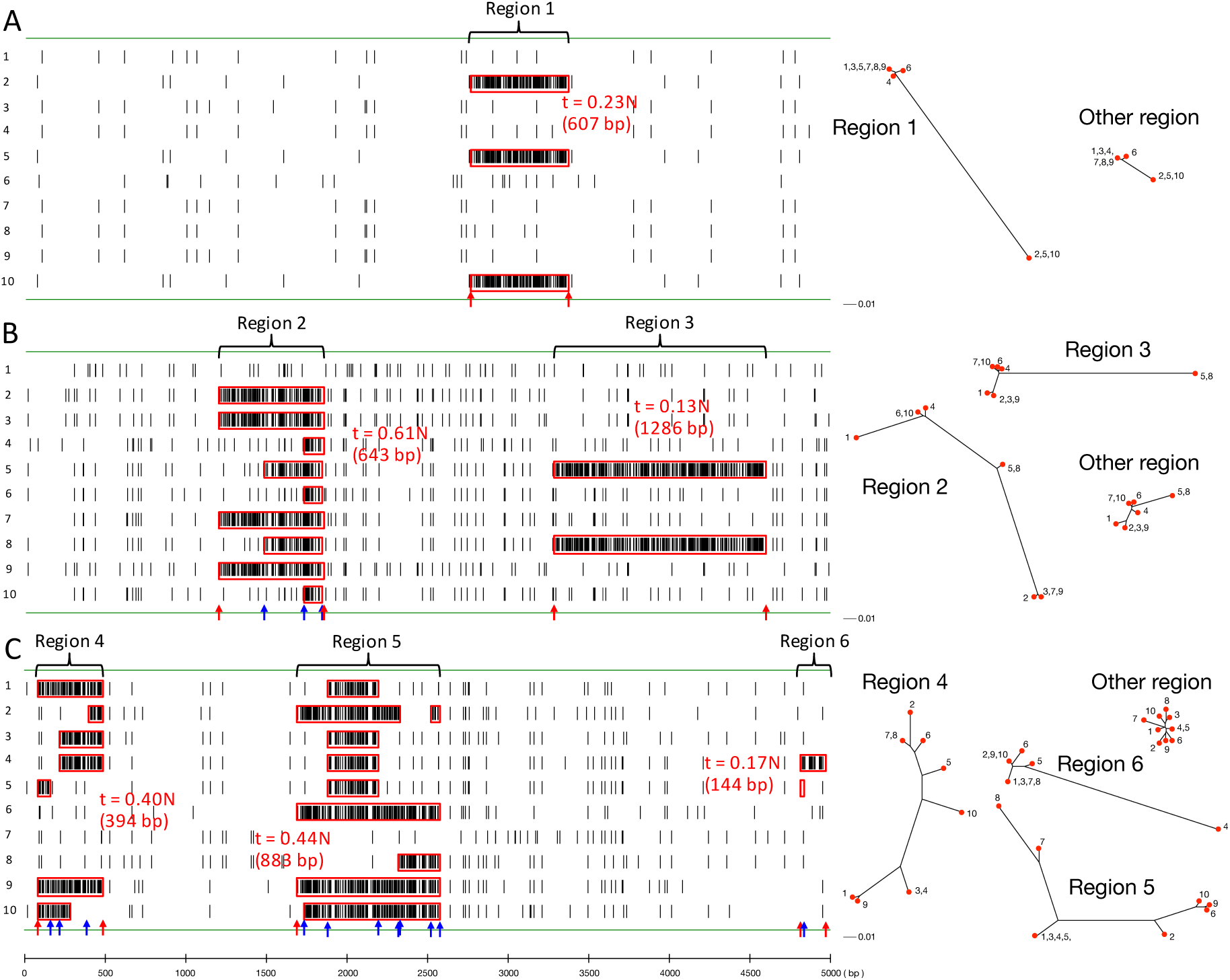
Typical patterns of SNPs with inter-specific recombination from external source with no intra-specific recombination (A; 2*Ng* = 0), with a moderate level of intra-specific recombination (B; 2*Ng* = 0.001), and with a high recombination rate (C; 2*Ng* = 0.005). Vertical bars indicate the locations of point mutations in the simulated region with *L* = 5000 bp. The regions that experienced inter-specific recombination are specified (Regions 1-6), and neighbor-jointing trees for these regions are shown in comparison with other regions with no inter-specific recombination. The breakpoints of inter-specific recombination events are presented by red allows, while blue ones exhibits intra-specific recombination events that fragmented the integrated foreign DNAs shown in red boxes.

**Figure 5.**
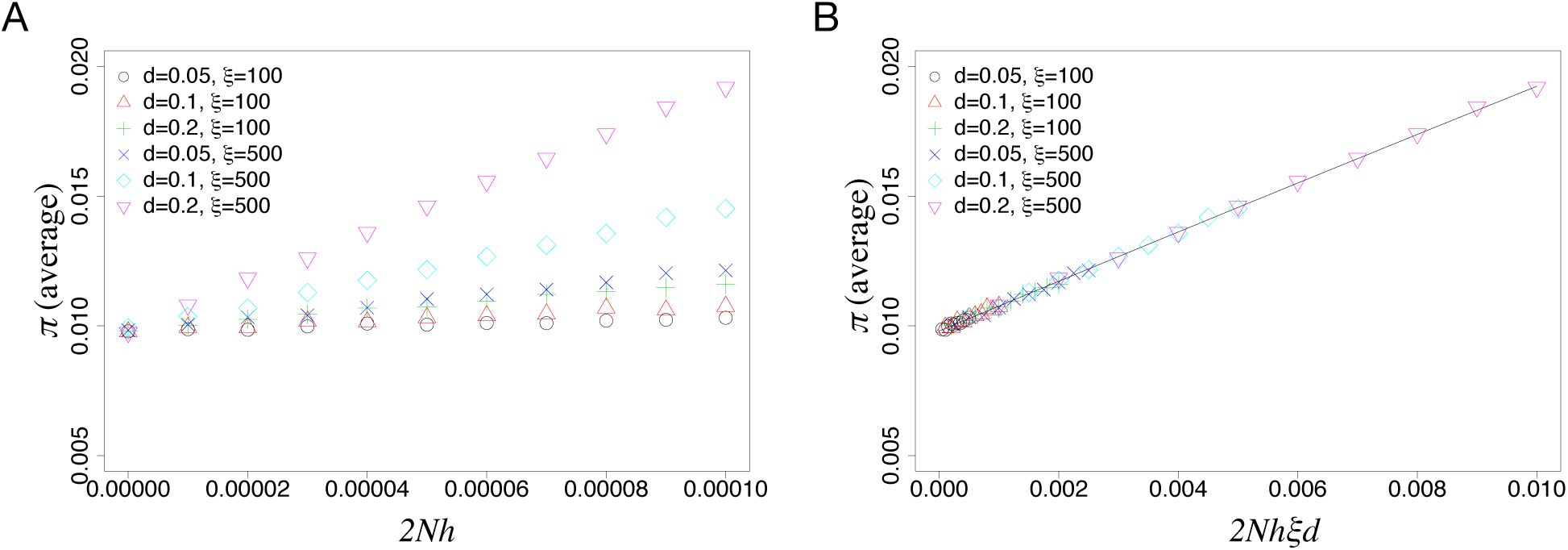
The effect of recombination from external source on the amount of polymorphism measure by *π* (A) *π* as a function of 2*Nh*. (B) *π* is in a clear linear correlation with 2*Nhξd* (Equation 8). The averages *π* over 10,000 runs of simulations with *n* = 15 and *L* = 10, 000 are shown.

In Figure 4B, a moderate level of recombination within species (intra-specific recombination) is introduced (2*N g* = 0.001). Two external DNA fragments are integrated (Regions 2 and 3). In Region 2, a 643 bp of foreign DNA was integrated *t* = 0.61N generations ago. It is important to notice that whereas individuals 2, 3, 7 and 9 have the entire fragment, only a part of the integrated fragment is observed in individuals 4, 5, 6, 8 and 10. This is due to intra-specific recombination that occurred after the integration; the integrated fragment was chopped into pieces and distributed into the population. By looking at the simulated ancestral recombination graph, we found three such intra-specific recombination events occurred (blue arrows the breakpoints). By contrast, in Region 3, due to its recent origin (*t* = 0.13N generations ago), no intra-specific recombination was involved so that the entire integrated region (1286 bp) remains intact in individuals 5 and 8, similar to Region 1 in Figure 4A.

With even a higher intra-specific recombination rate (2*Ng* = 0.005) in Figure 4C, fragmentation is more enhanced. There are three regions that experienced recombination from external source (Regions 4, 5 and 6), and all of them involved intra-specific recombination. An intriguing pattern is seen in Region 5, where only a part of the integrated flagrant is observed in the sample. The recombination occurred *t* = 0.44N generations ago. The actual length of the integrated foreign fragment was more than 883 bp, but none of the sampled ten individuals have the 5’ breakpoint. This process can be well understood with the cartoon in Figure 6, which illustrates the typical behaviors of population frequency of a foreign DNA integrated at time 0 with and without intra-specific recombination. With no intra-specific recombination, the entire integrated DNA can be vertically transmitted in the following generations (Figure 6A). By contrast, with intra-specific recombination, the integrated DNA is fragmented into various lengths (Figure 6B). As a consequence, more individuals have chances to have a part of the integrated DNA, but the length of the integrated DNA in each individual is on average short; some might lose the 5’ breakpoint and some might have only a short region in the middle. One potential caveat when interpreting data is that, when there was only one inter-specific recombination event, one might think multiple inter-specific events have incorporated foreign DNA independently. Indeed, when applied to one of our simulated data, **GENECONV** (Sawyer 1989), a commonly used software to detect gene conversion tracts, identifies a number of gene conversion tracts around the region that experienced a single time of inter-specific recombination, that incorporated a 2000 bp of foreign DNA at *t* = 0.94N (Figure 7). The tracts inferred by **GENECONV** are presented by purple lines, showing as if there is a hotspot of integration.

**Figure 6.**
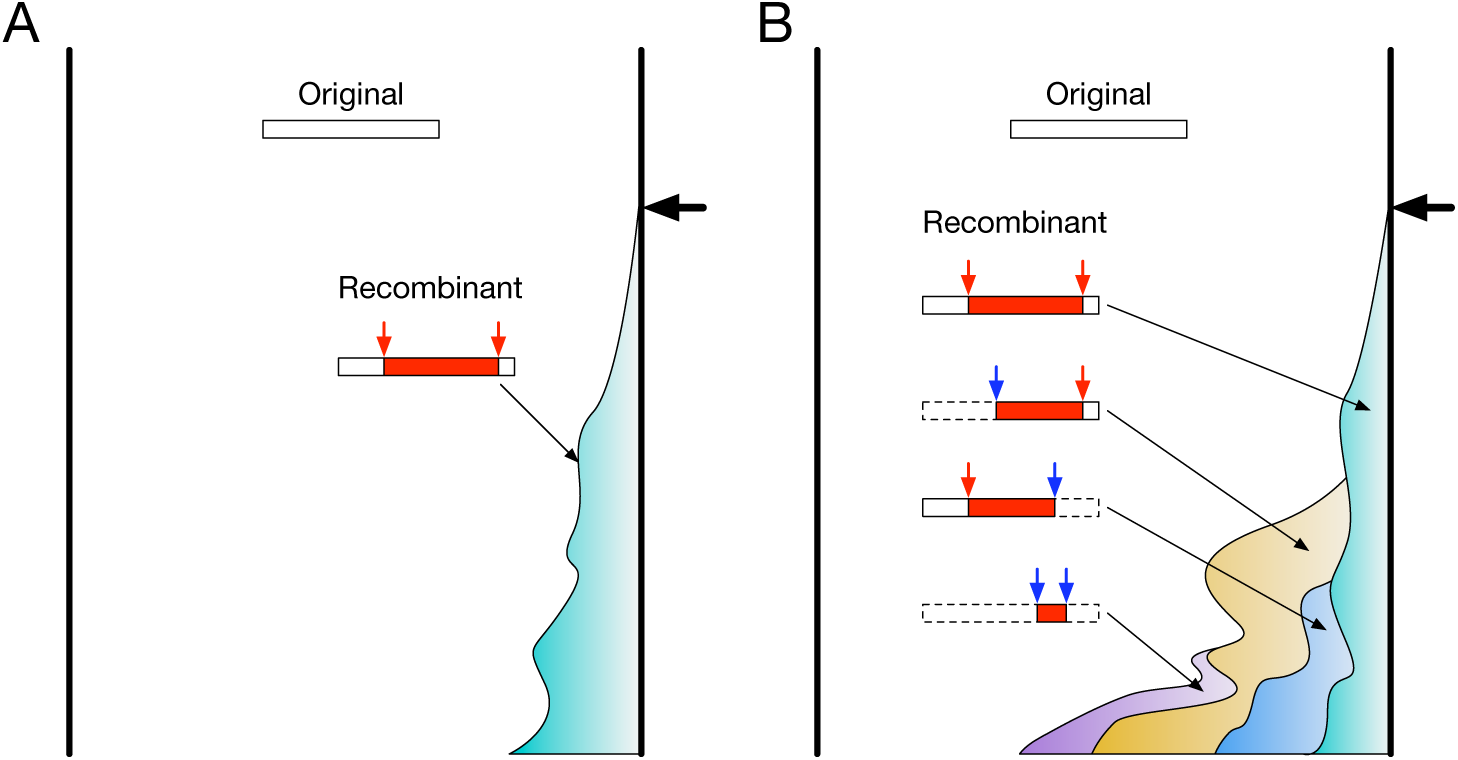
Cartoons of the typical behavior of population frequency of a foreign DNA, (A) without intra-specific recombination and (B) with intra-specific recombination. The time of the foreign DNA introduced into the population is denoted by a thick black arrow, producing a recombinant haplotype in which the integrated DNA is specified by a red box and arrows. (A) When there is no intra-specific recombination, the population consists of two haplotypes, the original and recombinant haplotype. (B) When intra-specific recombination is involved, the integrated DNA could be fragmented by recombination, thereby creating various kinds of recombinant haplotypes, each of which should have only a part of the integrated DNA. Additional breakpoints by intra-specific recombination are shown by blue arrows. In such a situation, the number of individuals having at least a part of part of the integrated DNA is much larger than the case with no recombination (A), while the length of integrated DNA is shorter.

**Figure 7.**
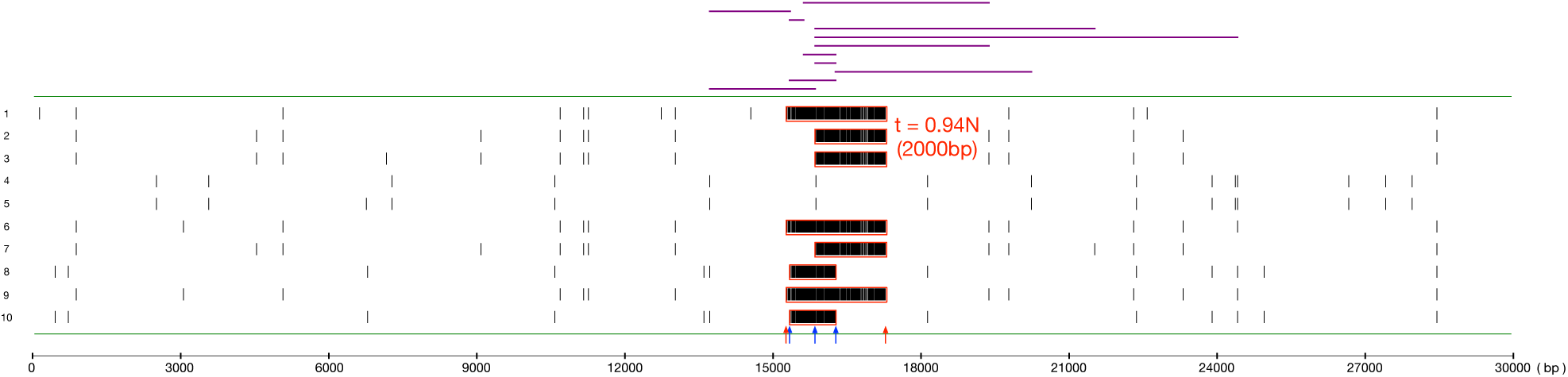
Application of **GENECONV** to a simulated region (*n* = 10, *L* = 30, 000 bp), in which a 2000 bp of foreign DNA was integrated 0.94*N* generation ago (red boxed), followed by three additional intra-specific recombinations that fragmented the foreign DNA. The breakpoints of inter-specific recombination events are presented by red allows, while blue ones exhibits intra-specific recombination events that fragmented the integrated foreign DNAs. Vertical bars indicate the locations of point mutations. **GENECONV** with the default setting identified 11 integrated tracts (purple horizontal bars), making it look as if there is a hotspot of integration.

Given this effect of inter-specific recombination, it is predicted that with increasing the rate of intra-specific recombination, (i) the number of individuals having foreign DNA increases and (ii) the length of foreign DNA decreases. This is quantitatively demonstrated by simulations (Figure 8). Figure 8 shows that the number of individuals that have at least a part of foreign DNA increase as the rate of intra-specific recombination (2*N g*) increases (Figure 8A), whereas the average length of each foreign DNA in the sample decreases (Figure 8B). This effect of intra-specific recombination (2*N g*) is larger when *ξ* is larger. These findings should be useful to improve the algorithms to identify genomic regions that underwent homologous recombination (Didelot and Falush 2007; Didelot *et al.* 2009; Ansari and Didelot 2014; Yahara *et al.* 2014).

**Figure 8.**
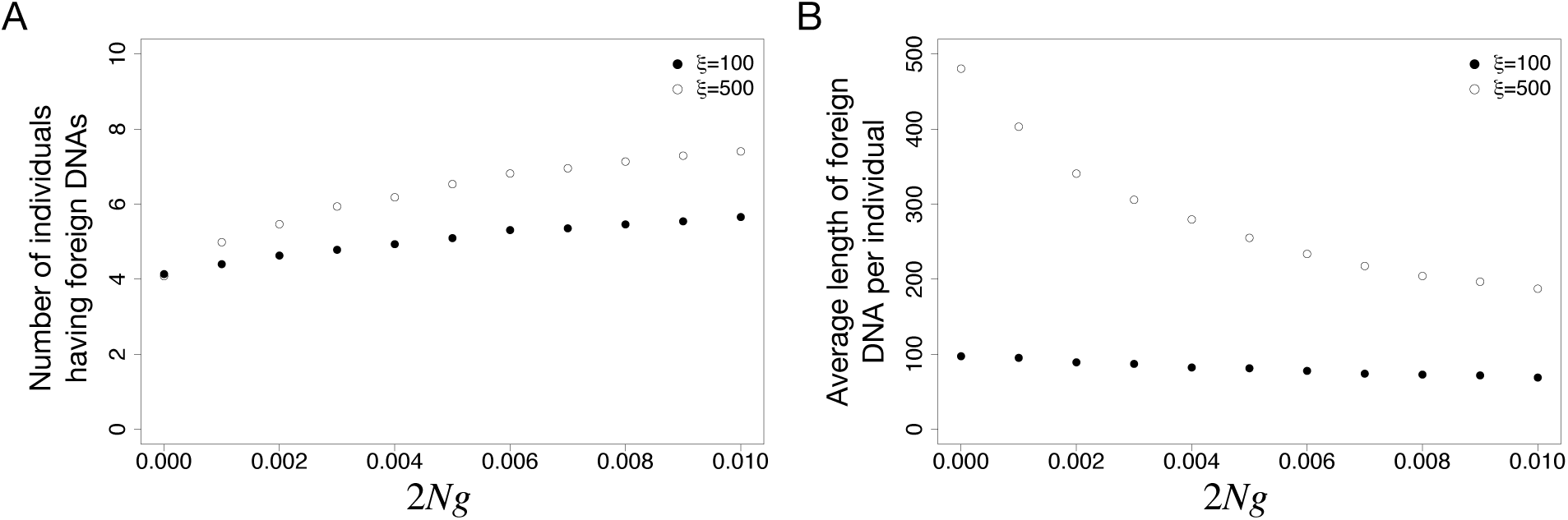
The effect of intra-specific recombination on (A) the number of individuals having foreign DNA and (B) average length of integrated foreign DNA, from 10,000 runs of simulations with *n* = 15 and *L* = 10, 000.

We thus demonstrated that the joint work of intra- and inter-specific recombination could create a complicated pattern of SNPs and it is needed to obtain full theoretical understanding of this for interpreting SNP data from prokaryotes. Given quite common homologous recombination from external source in prokaryotes and strong impact on the pattern of SNPs as we have shown here, we have to avoid a misleading interpretation of observed data due to recombination, potentially resulting in misevaluation of the relative contribution of demography and selection. We here developed a fast simulator for producing a number of realizations of SNPs with both intra- and inter-specific recombination. The software named **msPro** was developed based on Hudson’s commonly used software **ms (msPro** means **ms** for prokaryotes), and the input command and the form of output are very similar to **ms. msPro** can incorporate various forms of demographic history as **ms** does. **msPro** will be available upon request.

It should be noted that our simulator runs after specifying the density distribution of external DNA, *Q*_ext_. When there is no prior knowledge on the environmental DNA, it is difficult to set *Q*_ext_. Considering such a case, the default setting of **msPro** is given as follows. A first approximation is that the density distribution of tract length (*z*′) follow a geometric distribution *ξ*^−1^(1 − *ξ*^−1^)^*z*′−1^, regardless of divergence. According to empirical studies (Zawadzki and Cohan 1995; Linz *et al.* 2000), typical lengths of integrated DNA may be a few kb, so we assume *ξ* = 1000 bp. If we assume a uniform distribution of divergence in the external DNA in the environment, the density distribution of *d* (*i.e.,* divergence of successfully integrated DNA) simply follows the rate of successful integration, which may be approximated by an exponential distribution (see Fraser *et al.* 2007, and references therein), namely, *α* exp[−*αd*], where *α* is a parameter to specify the decay. According to Figure 1A in Fraser *et al.* (2007), *α* ~ 20 might fit the observed data from some bacterial species. Therefore, we set *Q*_ext_ (*d, z*′) = *α* exp[−*αd*] × *ξ*^− 1^(1 − *ξ*^−1^)^*z*′ − 1^.

## Acknowledgements

This work was supported in part by the Japan Society for the Promotion of Science (JSPS).

## Appendix

Consider a certain site of two samples and changes of their state in one generation backward in time. Let *P_t_* be their current diversity and *P*_*t*−1_ be that of before generation. Assuming that *de novo* mutation and recombination between species does not occur simultaneously and occur once at most, recursion of the state of their diversity under a finite two-states model can be written,

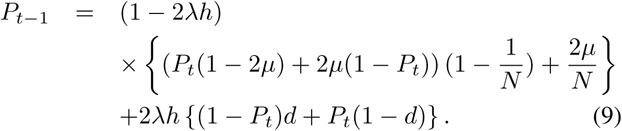

The expression in the first curly bracket means the case that the recombination does not occur, while the expression in the second curly bracket means the opposite case. At equilibrium (*i.e*., *P*_*t* − 1_ = *P_t_*), the recursion can be solved and then the states (denoted by *P*\*) is,

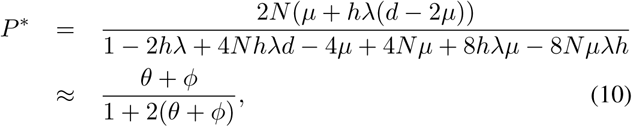

where *θ* = 2*N μ* and *ϕ* = 2*Nhλd,* and the terms with *h* or *μ* as factors are ignored. *P*\* is corresponding to the expectation of *π* in our framework.

